# A curated database reveals trends in single-cell transcriptomics

**DOI:** 10.1101/742304

**Authors:** Valentine Svensson, Eduardo da Veiga Beltrame, Lior Pachter

## Abstract

The more than 500 single-cell transcriptomics studies that have been published to date constitute a valuable and vast resource for biological discovery. While various “atlas” projects have collated some of the associated datasets, most questions related to specific tissue types, species, or other attributes of studies require identifying papers through manual and challenging literature search. To facilitate discovery with published single-cell transcriptomics data, we have assembled a near exhaustive, manually curated database of single-cell transcriptomics studies with key information: descriptions of the type of data and technologies used, along with descriptors of the biological systems studied. Additionally, the database contains summarized information about analysis in the papers, allowing for analysis of trends in the field. As an example, we show that the number of cell types identified in scRNA-seq studies is proportional to the number of cells analysed. The database is available at www.nxn.se/single-cell-studies/gui.

## Introduction

The availability of large numbers of comprehensive single-cell transcriptomics studies (Svensson, Vento-Tormo, and Teichmann 2018) is making possible the study of biological variation in unprecedented detail (Klein and Treutlein 2019). One interesting aspect of this “big data” biology consisting of a large set of measurements from many cells is that it can yield insights even after initial published analysis of individual datasets. Moreover, hundreds of datasets available, integration becomes a powerful tool for exploration. However, integration of diverse datasets requires standardization in how data is collected, shared, and curated (Stuart *et al.* 2019).

A number of “atlas” projects have been launched to address this problem and to assist researchers in focused domains. For example, The Human Cell Atlas portal aims to provide uniformly processed single-cell genomics data from all of the human body (Regev *et al.* 2017). JingleBells provides single-cell data, with a focus on immune cells (Ner-Gaon *et al.* 2017). The conquer database provides uniformly processed single-cell expression data to facilitate benchmarking of computational tools (Soneson and Robinson 2018). The PanglaoDB database provides single-cell RNA-seq count matrices from public sequencing data in the National Center for Biotechnology Information Sequence Read Archive (Franzén, Gan, and Björkegren 2019). The EMBL-EBI Single Cell Expression Atlas provides uniformly processed data from submissions to ArrayExpress. The Broad Institute offers a Single Cell Portal which can be used to share custom scRNAseq data. A database called scRNASeqDB provides links to a number of datasets from human scRNA-seq experiments (Cao *et al.* 2017). These efforts all aim to tackle different aspects of the considerable challenge of data management resulting from the extraordinary rapid adoption of single-cell genomics technologies.

We focus on a missing resource, namely a database of single cell transcriptomics studies rather than primary data. The compilation of such a database required us to read and manually curate large numbers of publications, which we indexed according to publication and study authors. Our database will allow researchers interested in specific tissues to rapidly identify relevant studies. Furthermore, by virtue of providing a comprehensive overview of the field, our database can highlight understudied tissues. Furthermore, the database will facilitate appropriate citation of previous work when performing follow-up experiments. The database tracks metadata applicable to most studies, such as the number of cell types identified, and protocols used. We show that these annotations enable analysis of trends in the field.

## Database structure

This database aims to provide a link between datasets from different tissues, pointers to data location, and relevant references. Together, these attributes make published data and results readily discoverable. A secondary goal is to annotate useful metadata associated with the primary studies.

The “*Single-cell studies database*” considers the analysis of many genes at once in single cells as a “single-cell transcriptomics” study. To allow for comprehensive coverage within a meaningful domain, the scope of the database was restricted in certain ways. For example, multicolor fluorescence flow cytometry and mass cytometry experiments were not included, even though both technologies can measure dozens of analytes per cell. The focus was restricted to datasets with the expression of more than a hundred genes measured in individual cells. Some targeted technologies measuring fewer genes such as osmFISH were also included when they could be directly related to higher throughput counterparts (Codeluppi *et al.* 2018; Shah *et al.* 2016; Wang *et al.* 2018).

The primary identifier of each entry in the database is the canonical digital object identifier (**DOI**) of a publication. Based on the DOI four entries are included using the CrossRef API: **Authors**, **Journal**, **Title** and **Date**. Additional fields are based on the contents of the publication and are manually annotated by investigating the text and supplementary material of the publication. If the study was deposited to the bioRxiv the **bioRxiv DOI** field indicates this. Attributes include:

**Reported cells total:**is the number of cells investigated in the study.
**Technique:** the technology or protocol used.
**Panel size:** the number of genes investigated when targeted technologies such as multiplexed smFISH were applied.
**Measurement:** the type of quantitative measurements performed (e.g. *RNA-seq, In Situ* or *Microarray*).
**Data location:** the public repository accession ID for the raw data.
**Organism:** the species of origin of cells examined in the study.
**Tissue:** the tissue type from which single cells were collected.
**Cell Source:** notes about the cells in the study.
**Contrasts:** the different experimental conditions studied, if any.
**Isolation:** the method used to produce the single cell suspension.
**Developmental stage:** the developmental stages or ages of the organisms cells were collected from.

Additionally, some fields are binary corresponding to a “Yes” or “No” entry. This is used for the following attributes:

**Cell clustering**: Did the study performed unsupervised clustering of cells (Islam *et al.* 2011)?
**Pseudotime**: were cellular trajectories inferred with pseudotime methods (Magwene, Lizardi, and Kim 2003)?
**RNA velocity**: was a vector field inferred from spliced and unspliced reads (La Manno *et al.* 2018)?
**PCA**: was a principal component analysis performed?
**tSNE**: was the t-Distributed Stochastic Neighbor Embedding algorithm used for visualization (Van der Maaten and Hinton 2008)?

Finally, the number of cell types or clusters identified in each study is recorded under **Number of reported cell types or clusters**. This is most commonly based on de novo clustering, but in some cases it is based on the number of distinct pre-sorted cell types.

While the manual curation of the data made possible description of numerous details from the papers in the database, some entries are missing due to difficulty in finding information. However, we believe the overall content of the database is substantial enough to serve as a good starting point for the community to contribute and fill in the gaps. We show that even with some missing annotation, the database in its current form makes possible analysis of trends in the field.

The database can be accessed via a graphical interface using Google Sheets at www.nxn.se/single-cell-studies/gui. This view allows searching on keywords and for browsing studies. Importantly, it also allows for the contribution of information to the database through comments on individual entries.

A version of the database in TSV (tab separated values) format can be downloaded from www.nxn.se/single-cell-studies/data.tsv. This enables researchers perform analyses using the data.

New studies can be submitted through a form located at www.nxn.se/single-cell-studies/submit. Submissions require a DOI. The form also allows for entry of additional metadata through optional fields. Claims in the submissions are spot checked to ensure they refer to the original text in the publication.

Every day a snapshot of the database is saved (in TSV format) using Google Cloud Functions, and all these snapshots are available in a public Google Storage bucket at gs: /single-cell-studies. An example snapshot is provided as Supplementary Table 1, which has data on 550 studies published between 2003 and August 17 2019.

## Results

The earliest single cell transcriptomics study recorded in the database was published in 2004. Since 2013, almost every month at least one study has been published. The rate of study publication has increased steadily, and in May, June, and July of 2019 there were over 30 single cell transcriptomics studies published per month **(Figure 1)**. In 2019 the median scRNA-seq study investigated approximately 14,000 cells **(Table 1)**.

**Figure 1.**
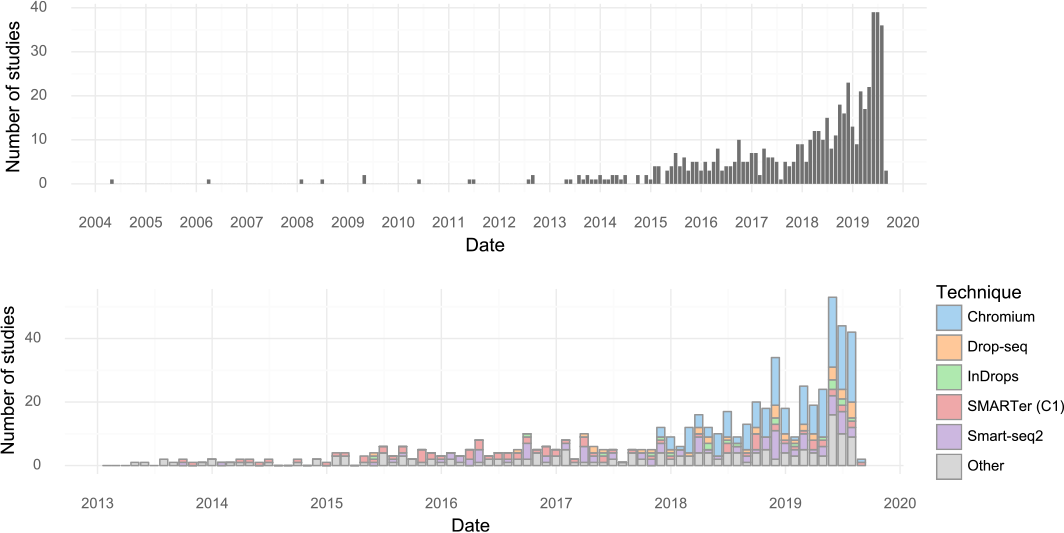
Studies over time. (**upper**) The number of single cell transcriptomics studies published per month. (**lower**) The number of scRNA-seq studies published per month stratified by method.

**Table 1.** Single cell study trends. (**left**) Number and size of single cell transcriptomics studies in 2019. (**middle**) Most common tissue investigated with single cell transcriptomics. (‘Culture’ refers to *in vitro* studies of cell lines). (**right**) Journals which have published most single cell transcriptomics studies. (‘bioRxiv’ means the study is so far only available on bioRxiv).

| Month | Studies | Median cells | Tissue | Studies | Journal | Studies |
| --- | --- | --- | --- | --- | --- | --- |
| Jan 2019 | 9 | 3,368 | Brain | 64 | bioRxiv | 63 |
| Feb 2019 | 21 | 11,175 | Culture | 47 | Nature | 50 |
| Mar 2019 | 16 | 11,452 | Blood | 16 | Cell | 49 |
| Apr 2019 | 21 | 17,725 | Heart | 16 | Cell Reports | 35 |
| May 2019 | 39 | 14,585 | Pancreas | 16 | Science | 34 |
| Jun 2019 | 39 | 15,000 | Embryo | 14 | Nature Communications | 29 |
| Jul 2019 | 36 | 13,966 | Lung | 12 | Genome Biology | 19 |

Individual studies have increased in scale over time, and every few months a new study is released that breaks the previous record in terms of number of cells assayed. During the first half of 2019 approximately 200,000 cells were added to the pool of public data every month **(Figure 2)**.

**Figure 2.**
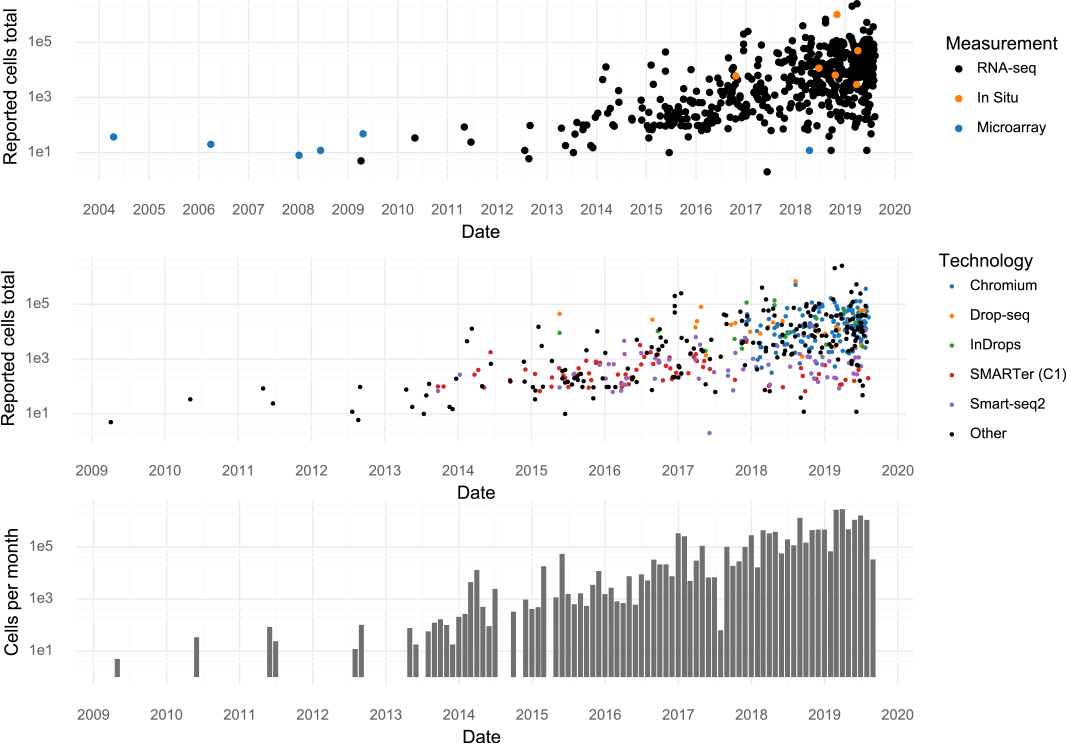
Scale of experiments and data over time. (**Upper**): The number of cells measured in a study, stratified by the measurement method. (**Middle**): The number of cells measured in scRNA-seq experiments, stratified by scRNA-seq protocol. (**Lower**): The aggregate number of cells measured per month.

Many tissues have been investigated by single cell transcriptomics methods, but the brain is the most popular with 65 associated cita tions out of 550 **(Table 1)**. Another trend observed from this database is that authors of single cell transcriptomics papers are increasingly making use of the bioRxiv preprint server. In total 145 of 555 studies were deposited to bioRxiv (26%). The fraction is now about 41% in a given month **(Figure 3)**. Single cell studies are published in many different journals, with Nature and Cell having published the most **(Table 1)**. The increasing popularity of these kinds of studies means the field, as measured by number of active authors, has grown. There have been 5,823 distinct authors of single cell transcriptomics studies.

**Figure 3.**
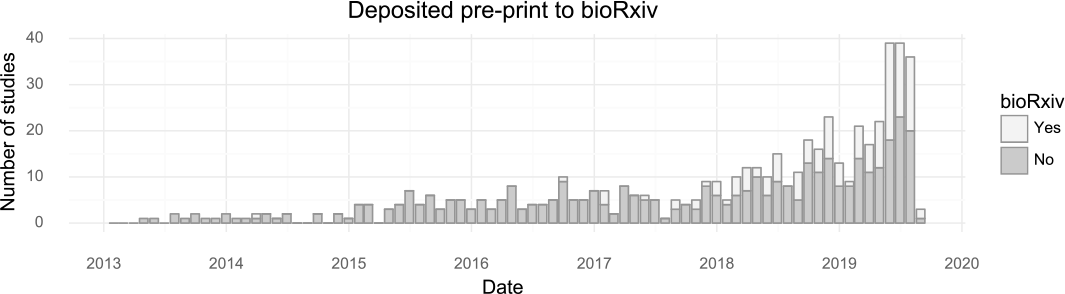
Pre-print usage over time. The number of studies published in a given month stratified by whether they at some point were deposited to bioRxiv. (Including studies currently only available on bioRxiv).

By tracking what forms of analyses are performed with single cell transcriptomics data, it is possible to learn something about what the community as a whole is aiming to learn from the assays. The most common application is to survey molecular “cell types” by clustering cells based on gene expression. Almost every study performs clustering (87%). The t-SNE visualization method became nearly universally applied after its first use for single-cell analysis in 2015, although the fraction of studies per month using it has decreased slightly in the last year, possibly due to the introduction of UMAP (McInnes and Healy 2018). “Pseudotime” is less frequently examined but is still very popular with about half of published studies investigating pseudotime trajectories **(Figure 4)**.

**Figure 4.**
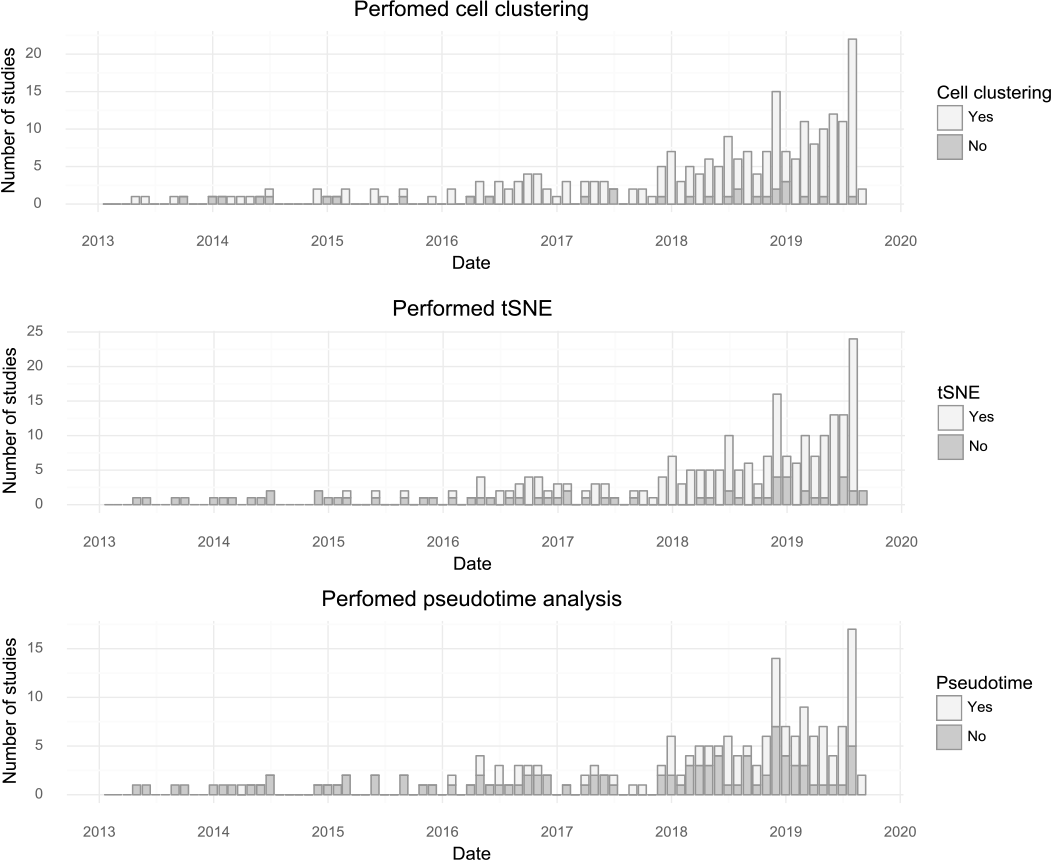
Popularity of forms of analysis over time. (**Top**) The number of studies doing clustering per month. (**Middle**) The number of studies using tSNE per month. (**Bottom**) The number of studies doing pseudotime analysis per month.

Since *de novo* clustering and cell type discovery is almost always performed, we annotated the number of clusters of cells identified in the studies. This revealed a high correlation between cell type numbers and the number of cells investigated. For small to medium sized studies on average one cell type is identified per 100 cells assayed. For large studies with hundreds of thousands of cells, the rate is closer to one cell cell type per 1,000 cells assayed **(Figure 5)**.

**Figure 5.**
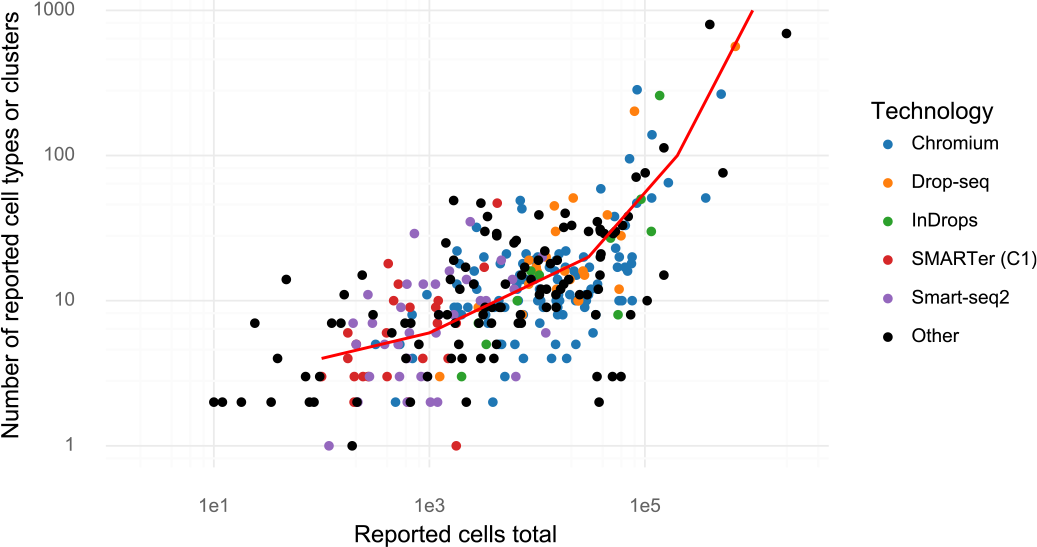
Cluster and cell numbers. The number of cells studied vs the number of clusters or cell types reported in a study.

The curated database described here is hosted at www.nxn.se/single-cell-studies. It has been designed for easy access to the underlying data and for in depth analysis in Python or R. The database was designed to facilitate access to published single-cell research, so that for example a researcher can find all single cell studies of the pancreas to explore the results and analyze public data. We found that analysis of other aspects of the studies described in the papers, namely attributes such as type of protocol, number of cells, or the number of clusters identified, revealed interesting trends in the field. We believe that our finding that the number of clusters identified is directly proportional to the number of cells analysed merits some scrutiny in light of the biological significance that is frequently associated with the number of clusters detected.

The database is also designed to enable contributions by the community via a mechanism for suggesting additions, adding data, and for commenting. Forms for these functions are hosted at www.nxn.se/single-cell-studies/gui and www.nxn.se/single-cell-studies/submit.

## Acknowledgements

We thank Carlos Talavera-López for helpful feedback on the manuscript. Cloud infrastructure was funded through the Google Cloud Platform research credits program. The work was partly funded by NIH U19MH114830.

